# The protective effect of NGF mutant CHF6467 in SH SY5Y cells exposed to Aβ(1–42): a pharmacometabolomic approach

**DOI:** 10.64898/2026.10.06.753538

**Authors:** Enza Napolitano, Carmen Marino, Manuela Grimaldi, Michela Buonocore, Angelo Santoro, Gaetano Barbato, Anna Maria D’Ursi

## Abstract

Nerve Growth Factor (NGF) supports neuronal survival, plasticity, and growth by binding to p75NTR and TrkA receptors. There is a strong link between NGF and the cholinergic hippocampus, and changes in NGF pathways are observed in Alzheimer’s disease (AD). Research shows NGF’s neuroprotective and regenerative effects on cholinergic neurons, with positive results in AD models and clinical studies. However, NGF therapy faces challenges: limited blood–brain barrier crossing and dose-dependent pain. To address pain, a mutated NGF called CHF6467 was developed. Thanks to the R100E mutation, it prevents binding to p75NTR while maintaining TrkA affinity, which helps reduce pain side effects. Understanding CHF6467’s mechanism remains a challenge. In this context, untargeted metabolomics is effective for uncovering unforeseen responses, making it suitable for assessing CHF6467’s overall effects. Building on our previous research and recognizing the importance of amyloid-β peptide as a critical pathological marker of AD, we used NMR-metabolomics to study the effect of CHF6467 on SH-SY5Y cells. The study examined metabolomic profiles of cells treated with CHF6467, with and without Aβ(1-42), using NMR. Results show CHF6467 influences the metabolic changes caused by Aβ(1-42), inducing biochemical effects consistent with the promotion of phospholipid precursors, reduced glutamate excitotoxicity, and restored energy metabolism.

## Introduction

Nerve Growth Factor (NGF) is a key neurotrophic factor that supports neuronal survival, plasticity, and growth by binding to two receptors: the low-affinity p75 neurotrophin receptor (p75NTR) and the high-affinity tropomyosin receptor kinase A (TrkA). Both receptors are expressed in basal forebrain cholinergic neurons and are known to innervate the hippocampus. ^1^

Alterations in the NGF metabolic pathway have been observed, particularly in basal forebrain cholinergic neurons of patients with Alzheimer’s disease (AD). Among the various hypotheses for AD development, the NGF hypothesis is prominent. ^2^ It is supported by the well-known, tightly regulated connection between cholinergic neurons and NGF signaling, indicating that disruptions in NGF availability or signaling could lead to the degeneration of these neurons. Accordingly, cognitive deficits in AD are linked to cholinergic neuron loss, and the basal forebrain cholinergic system is the only brain region that relies on NGF during postnatal development. Additionally, it has been demonstrated that NGF increases choline acetyltransferase activity in this brain region when injected into the cerebrospinal fluid and protects basal forebrain cholinergic neurons following axonal damage to fibers projecting from the hippocampus. ^3, 4^

Multiple dysfunctions in NGF signaling, transport, maturation, and degradation are characteristic of AD brains. ^5^

In basal forebrain tissue from AD patients, while cortical NGF mRNA levels often remain normal, the immature growth factor proNGF is increased.^6^ Enzymes necessary for proNGF maturation, such as tPA, plasminogen, and plasmin, are significantly reduced in postmortem AD brains. Additionally, MMP-9 activity, the main NGF-degrading protease, is notably higher compared to controls. ^7^ These combined effects result in lower levels of active mature NGF and a buildup of inactive proNGF. Research has extensively explored NGF’s neuroprotective and regenerative effects on cholinergic neurons, showing promising results in animal models and clinical trials. ^8, 9^ However, despite these benefits, NGF therapy faces two major challenges: poor ability to cross the blood–brain barrier (BBB) and the risk of dose-related pain as a side effect. ^10, 11^

To overcome pharmacokinetic limitations and boost NGF concentration in the brain, researchers have explored advanced delivery systems, including gene- and cell-based therapies, as well as NGF released from polymer-encapsulated cells. ^8, 12^ As regards the second limitation, a mutated NGF variant, CHF6467, has been developed. The CHF6467 sequence contains two mutations relative to wild-type NGF: P61S and R100E. The R100E mutation selectively disrupts NGF binding to the p75NTR receptor while maintaining its affinity for TrkA. This improves NGF’s tolerability in clinical use by reducing pain side effects. The P61S mutation was added solely as a marker for the recombinant form, enabling CHF6467 to be distinguished from wild-type NGF. ^13^

Previous studies have already examined the behavior of the CHF6467 mutant in both in vitro and in vivo models. Capsoni *et al.* showed that this mutant selectively activates the beneficial downstream pathways triggered by TrkA, while avoiding those involved in pain transmission. Thanks to this signaling profile, CHF6467 supports neuronal survival and differentiation without engaging nociceptive mechanisms. In animal models, such selective signaling prevents learning and memory deficits and protects against neurodegeneration. ^14^ The same authors later reported that the NGF mutant also modulates inflammatory mediators released by glial cells, including soluble TNFα receptor II and the chemokine CXCL12.^15^

Given the established role of NGF in neuroprotection and the emerging clinical potential of its mutant, CHF6467, elucidating its mechanism of action (MoA) using broader, less-restricted approaches represents a valuable opportunity in medicinal chemistry.

In this context, metabolomics is increasingly recognized as a valuable tool for MoA investigation, offering an integrated view of the biochemical perturbations triggered by treatment. ^16–21^

Metabolomics is the latest and most promising omics field that systematically analyzes metabolites, enabling both qualitative and quantitative identification to investigate the effects of disease, external stimuli, or treatments, as well as genetic modifications. In its untargeted form, metabolomics has proven to be a powerful and effective method, not only in discovering new disease biomarkers but also in understanding mechanisms of action, identifying drug targets, and uncovering potential off-target effects. Nuclear Magnetic Resonance (NMR) is the preferred technique in metabolomics when an untargeted approach is desired.^22, 23^ Untargeted NMR metabolomics has proven effective in studying MoA of already approved or new potential drugs, providing valuable insights into their biological effects.^17, 24–28^ Within this framework, we previously used this method to investigate the impact of the alkaloid nicotine in a simple cellular model, clarifying its role in stimulating neurons and offering protection against amyloid toxicity. ^29, 30^ These recent applications clearly show that untargeted metabolomics provides an unbiased approach to investigate MoA, ensuring thorough, methodologically sound analyses without relying on limiting assumptions.

Building on our previous research and recognizing the importance of amyloid β-peptide as a critical pathological marker of AD ,^30–32^ we used an NMR metabolomic approach to study how CHF6467 affects SH-SY5Y cells exposed to Aβ(1-42) peptide.

The SH-SY5Y neuroblastoma cell line is commonly used as an in vitro model in neurodegenerative disease research, helping minimize confounding factors and address ethical issues associated with in vivo models. ^31, 33, 34^ Although immortalized cell lines have inherent limitations, SH-SY5Y cells offer advantages such as genetic stability, reproducibility, and scalability. ^35, 36^

Based on these considerations, our study aimed to analyze the metabolomic profile of SH-SY5Y cells treated with CHF6467, both in the presence and absence of Aβ(1-42) peptide, using NMR spectroscopy. Our metabolomic analysis suggests that CHF6467 can effectively modulate the metabolic alterations induced by Aβ(1-42). The approach employed—combining the broad perspective of the untargeted method with the integration of endometabolites and exometabolites data—provides valuable insights into how CHF6467 acts. This improves our understanding of how the promising drug works, utilizing a powerful and emerging methodology to analyze MoA.

## Results

### 1. CHF6467 protects SH-SY5Y cells against Aβ (1-42) toxicity inducing a metabolic reprogramming

To assess CHF6467’s protective role on SH-SY5Y neuroblastoma cells, a cell viability assay was conducted. Results confirmed CHF6467’s protective effect against Aβ(1–42)-induced toxicity. As shown in **Figure 1**, SH-SY5Y cells treated with Aβ(1–42) (50 μM) had a survival rate of 50.12 ± 3.91%. Co-treatment with CHF6467 at 100 ng/mL and 200 ng/mL improved cell survival to 71.1 ± 0.18% and 75.56 ± 5.14%, respectively.

**Figure 1.**
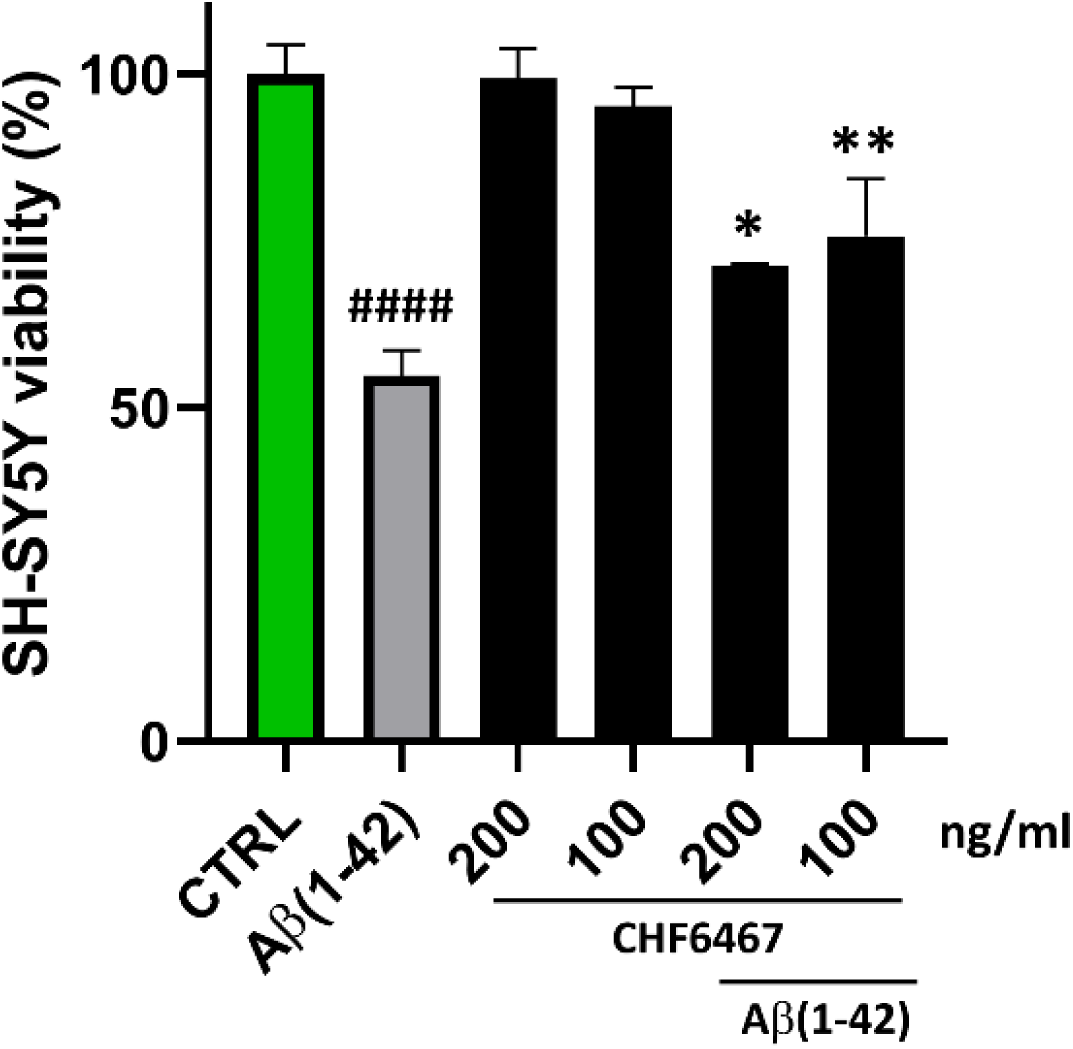
The neuroprotective effect of CHF6467against cytotoxicity induced by Aβ (1-42). Cell viability was assessed using the CCK-8 assay. SH-SY5Y cells were pretreated with CHF6467 at 100 ng/mL and 200 ng/mL for 24 hours, then exposed to 50 μM recombinant monomer of Aβ (1-42) for an additional 48 hours. Changes in viability were expressed as the percentage of living cells in treated cultures relative to untreated controls (CTRL). The results, presented as mean ± SD from three independent experiments, show significant differences: #### indicates p < 0.0001 versus Ctrl; * and ** denote p < 0.05 and p < 0.01, respectively, versus Aβ (1-42).

To explore how CHF6467 protects SH-SY5Y cells from Aβ(1–42) toxicity, we performed a combined metabolomic analysis of both intracellular and extracellular compartments. For the metabolomic study, a subtoxic 5μM concentration of Aβ(1-42), previously used in similar experiments, was selected. ^32, 37^ This choice ensures that metabolomic alterations induced by the pathological insult can be detected while remaining well below the EC_50_, whose impact on cell viability would otherwise compromise the reliability of metabolomic analysis. With regard to CHF6467, the lowest concentration (100 ng/mL) was selected for this exploratory study, in accordance with previous work examining the protective effects of NGF in cellular systems. ^38, 39^ ^1^H NOESY NMR spectra were acquired using cell extracts and medium samples to provide insights into the endometabolome and exometabolome, respectively.^40^ The data on the identification and quantification of metabolites were collected in a combined data matrix and subjected to multivariate statistical analysis.

Specifically, we used sparse Partial Least Squares Discriminant Analysis (sPLS-DA) to compare the metabolomic profiles of three cell groups: (i) cells treated with CHF6467 followed by Aβ(1–42) incubation (Abeta_CHF6467); (ii) cells incubated with Aβ(1–42) alone (Abeta); and (iii) untreated control cells (CTRL).

The score scatter plot shows that the Abeta_CHF6467 group is clearly separated from the Abeta group—representing the disease condition—but partially overlaps with the control group (**Figure 2A**). This suggests that the metabolomic profiles of both endo- and exometabolites in co-treated cells are similar to those in the control group, indicating that CHF6467 can restore the metabolomic profile to a healthy state.

**Figure 2.**
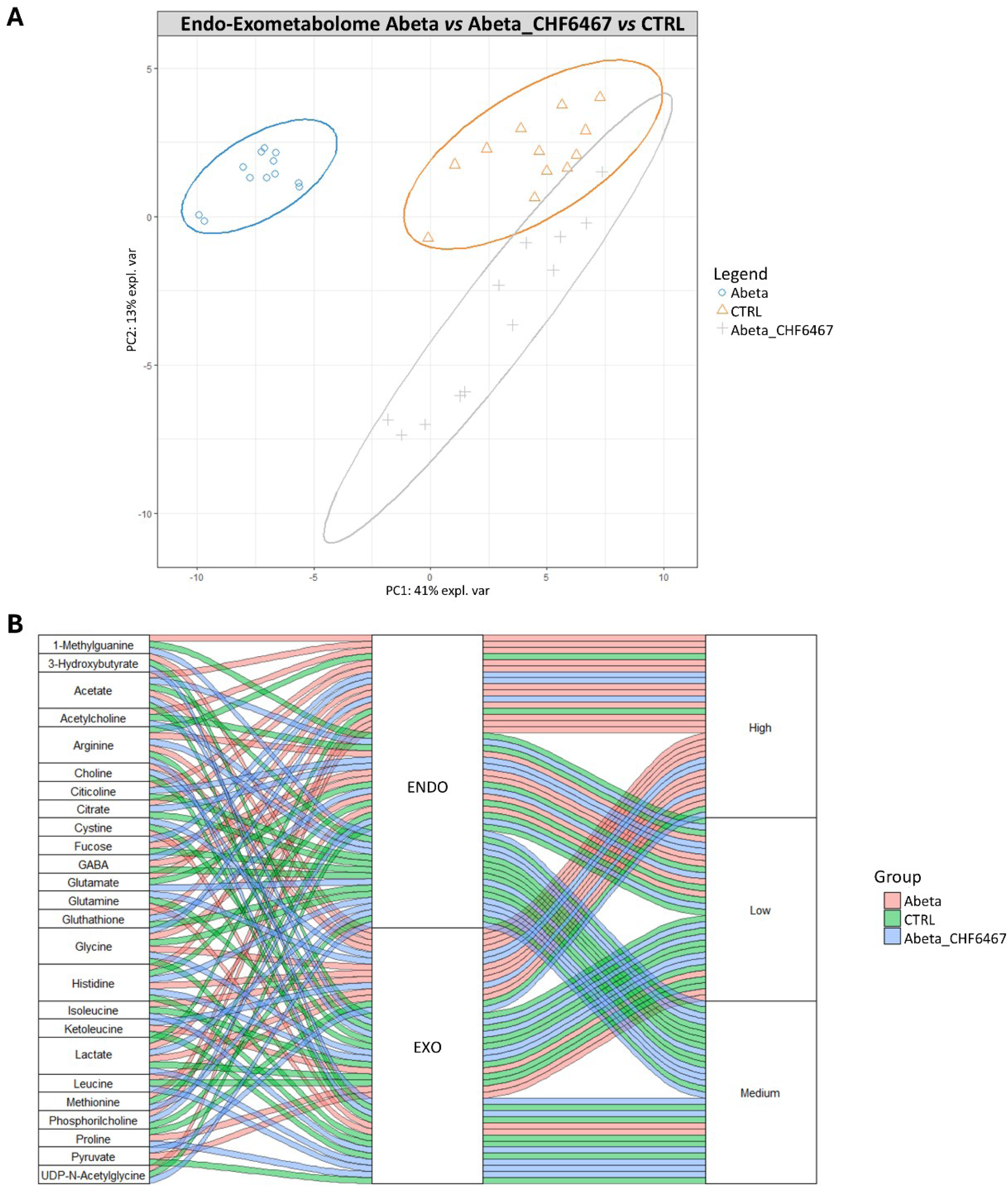
(**A**) sPLS-DA score scatter plots illustrate the combined endo- and exometabolome matrices of SH-SY5Y cells treated with Aβ (1-42) in blue (Abeta), cells pretreated with CHF6467 prior to Aβ (1-42) in grey (CHF6467_Abeta), and control cells in orange (CTRL). Cluster analyses are shown in a Cartesian space defined by principal components PC1: 41% and PC2: 13%. Cross-validation (CV) analysis assessed sPLS-DA, with CV tests following the statistical protocol showing significant cluster separation, with accuracy values of 0.63 and 0.86 on PC1 and PC2, and positive Q2 indices of 0.65 and 0.90. (**B**) Alluvial plot displaying metabolites that discriminate clusters analyzed via sPLS-DA and classified based on VIP values. The first column lists the discriminating metabolites, the second the cellular compartment, and the third the direction of concentration change. Max, Medium, and Min indicate the maximum, intermediate, and minimum concentrations across the three clusters. Lines connect each metabolite to its respective cell compartment, with changes in concentration color-coded: pink for metabolites in Aβ (1-42)-incubated cells, blue for cells pretreated with CHF6467 before Aβ (1-42), and green for control cells.

The model’s validity was confirmed through cross-validation, with Q2 values of 0.65 and 0.90 for PC1 and PC2, respectively. Additional validation via ROC curve analysis (**Figure S1**) supports these findings, indicating that CHF6467 significantly helps reverse the metabolic changes induced by Aβ(1–42).

The alluvial plot (**Figure 2B**) illustrates the metabolites most influential to this clustering, emphasizing compounds with Variable Importance in Projection (VIP) scores above 1 that show different levels across intra- and extracellular compartments under the three experimental conditions. It demonstrates that cells pretreated with CHF6467 before incubation with Aβ (1-42) exhibit low intracellular levels of 1-methylguanine, acetylcholine, glutamate, methionine, and UDP-N-acetylglycine, but high levels of fucose and GABA. Conversely, the exometabolome of this group exhibits the highest levels of glutamine, ketoleucine, and pyruvate, while showing lower levels of 3- hydroxybutyrate, isoleucine, leucine, and proline. Our analysis underscores the importance of metabolite exchange between the two cellular compartments, with notable changes observed in the concentrations of acetate, arginine, glycine, histidine, and lactate in both the endo- and exometabolomes.

**Figure 3** shows heatmaps of the average concentrations of intracellular and extracellular metabolites across three experimental cell groups. The data indicate that Aβ(1–42)-treated neuroblastoma cells in the presence of CHF6467 have a metabolomic profile very similar to that of the untreated control cells in both the endometabolome and exometabolome, thus indicating that CHF6467 treatment can alter the metabolic state of cells exposed to Aβ(1–42), bringing it closer to that of untreated cells.

**Figure 3.**
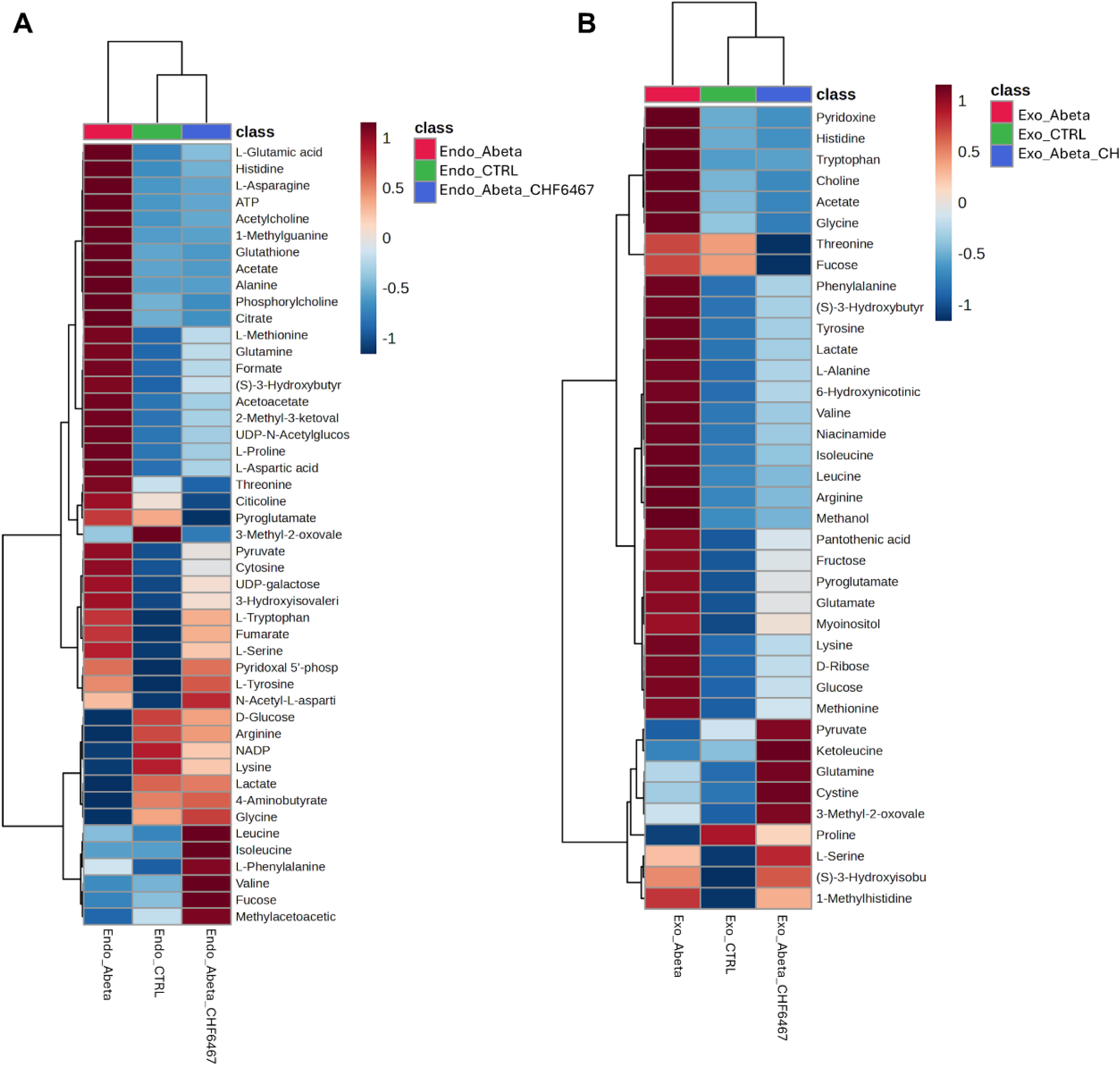
Heatmap showing altered metabolites in endo (**A**) and exometabolome (**B**). Each section’s color indicates the metabolite’s concentration, derived from a normalized concentration matrix (red for upregulated, blue for downregulated).

**Table 1** displays the results of the Pathway Enrichment Analysis, which interprets the NMR-based metabolomic data to identify biochemical pathways affected by CHF6467 in the presence of Aβ(1– 42). The analysis highlights changes in several amino acid-related pathways, such as glutamate, alanine, methionine, betaine, glycine, serine, aspartate, arginine, and proline metabolism. Additionally, lipid-related processes—including carnitine synthesis, phospholipid and phosphatidylcholine biosynthesis, and sphingolipid metabolism—are also impacted. Sugar metabolism pathways, including amino sugar metabolism, gluconeogenesis, the glucose–alanine cycle, and glycolysis, are significantly modulated. Lastly, the analysis reveals alterations in spermidine and spermine metabolism, as well as in glutathione metabolism.

**Table 1.**
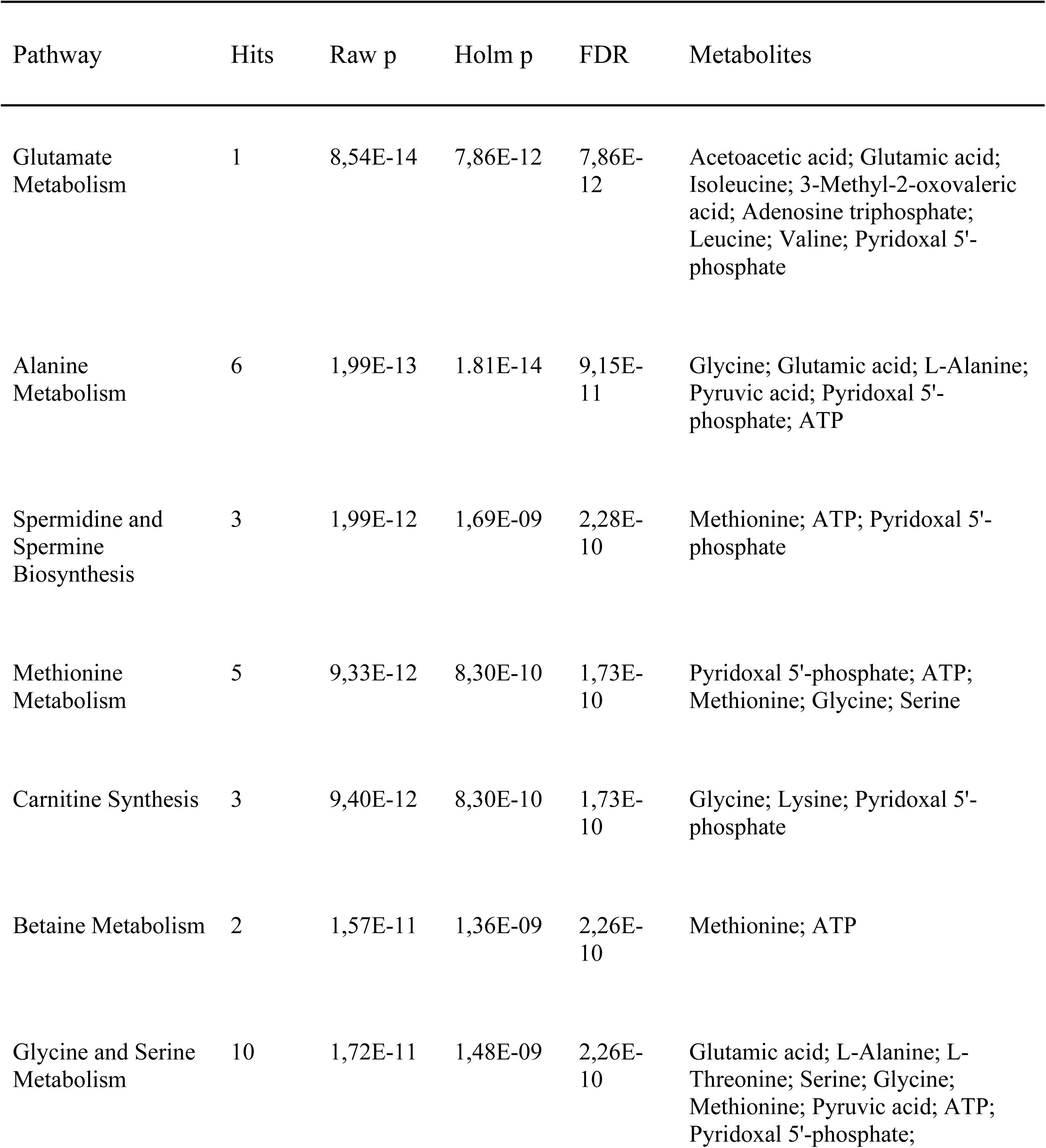
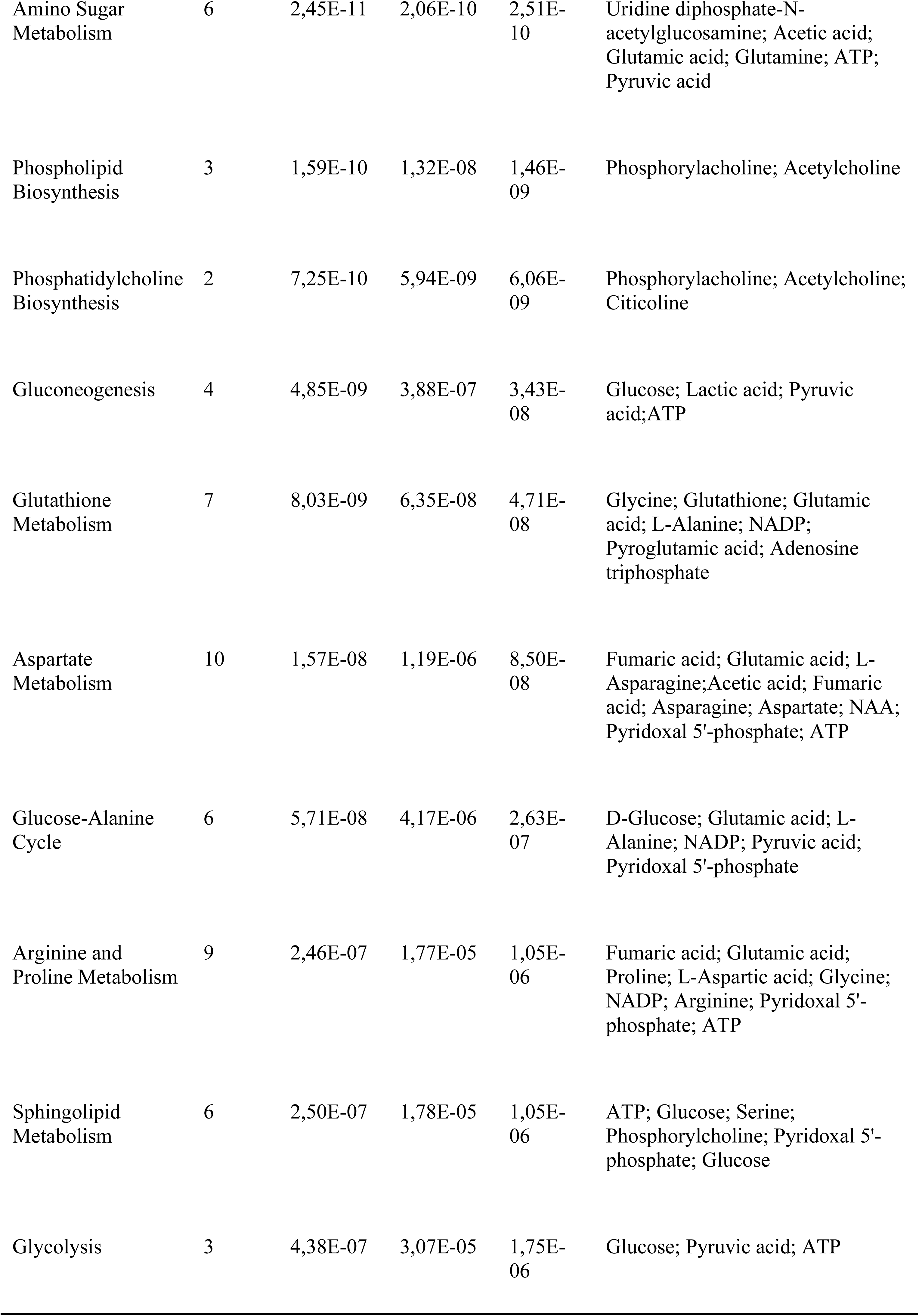
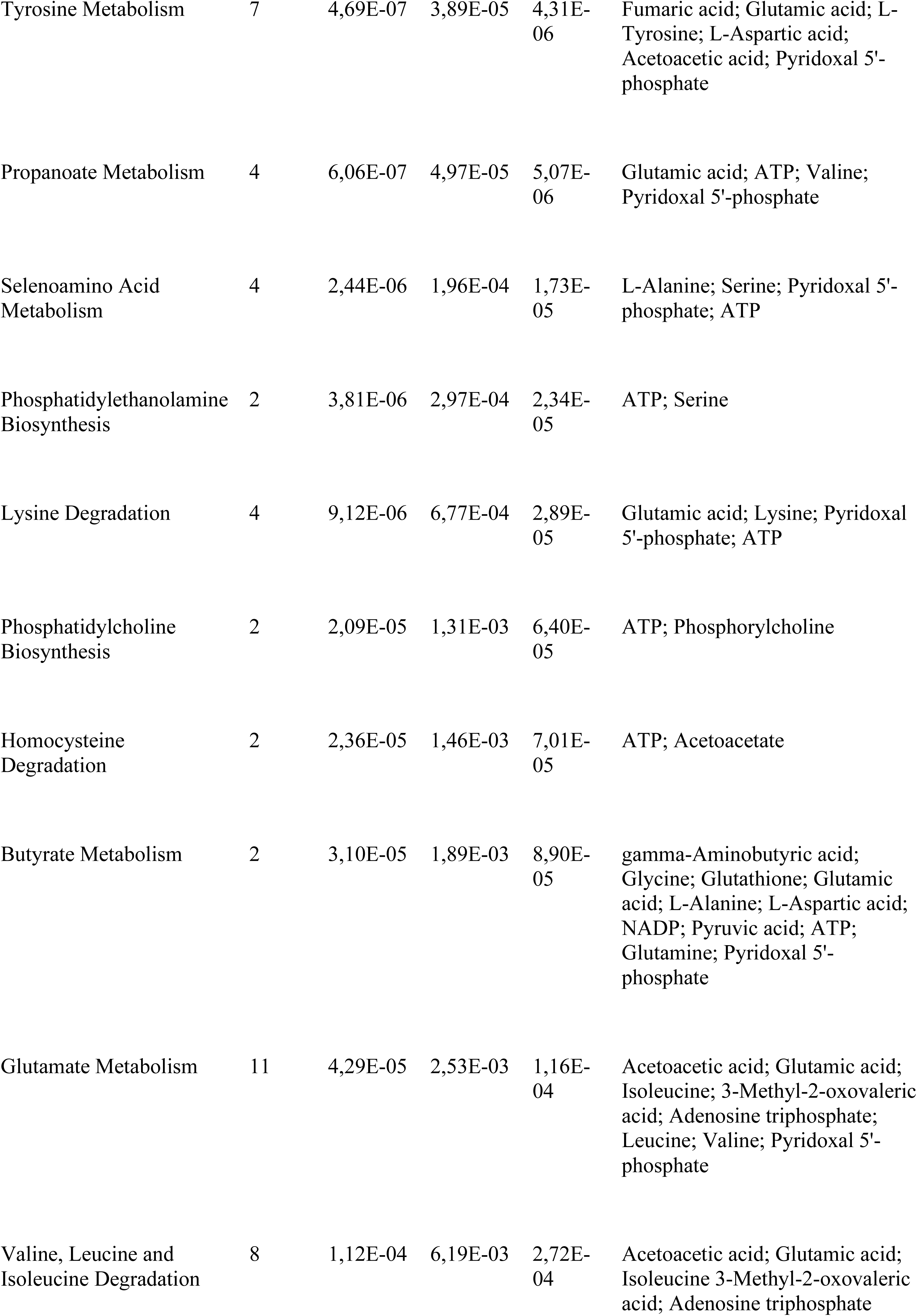
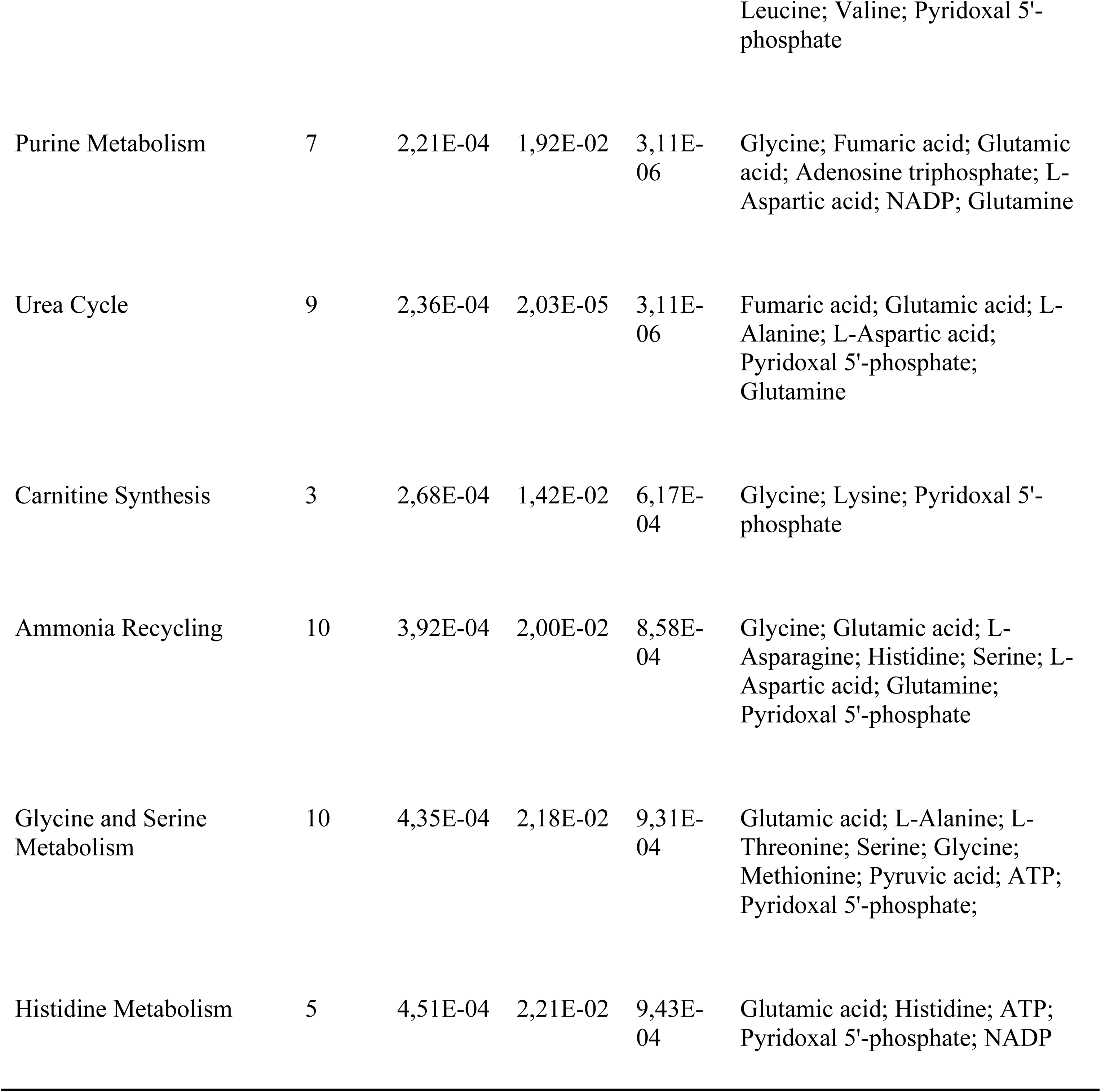
Enrichment pathway analysis: pathways distinguishing Aβ(1–42)-treated cells from cells exposed to both CHF6467 and Aβ(1–42). The table reports the raw p-value, Holm p, FDR value, and involved metabolites for each pathway. Only pathways with more than one hit and both raw and Holm-adjusted p-values below 0.05 are considered significant.

#### 3.3.3 CHF6467’s metabolic effects

To detect early biochemical changes that may explain CHF6467’s protective role against Aβ(1-42) toxicity, we examined the metabolomic profiles of SH-SY5Y neuroblastoma cells treated with CHF6467 (100 ng/mL for 24 hours).

We compared these profiles with untreated controls using sPLS-DA (**Figure 4A**). The score plot shows a distinct metabolic shift caused by CHF6467, impacting both intracellular and extracellular compartments. The model’s performance was validated through cross-validation, with Q2 indices of 0.94 for PC1 and 0.85 for PC2. Further confirmation of the model’s robustness came from ROC curve analysis (**Figure S2**).

**Figure 4.**
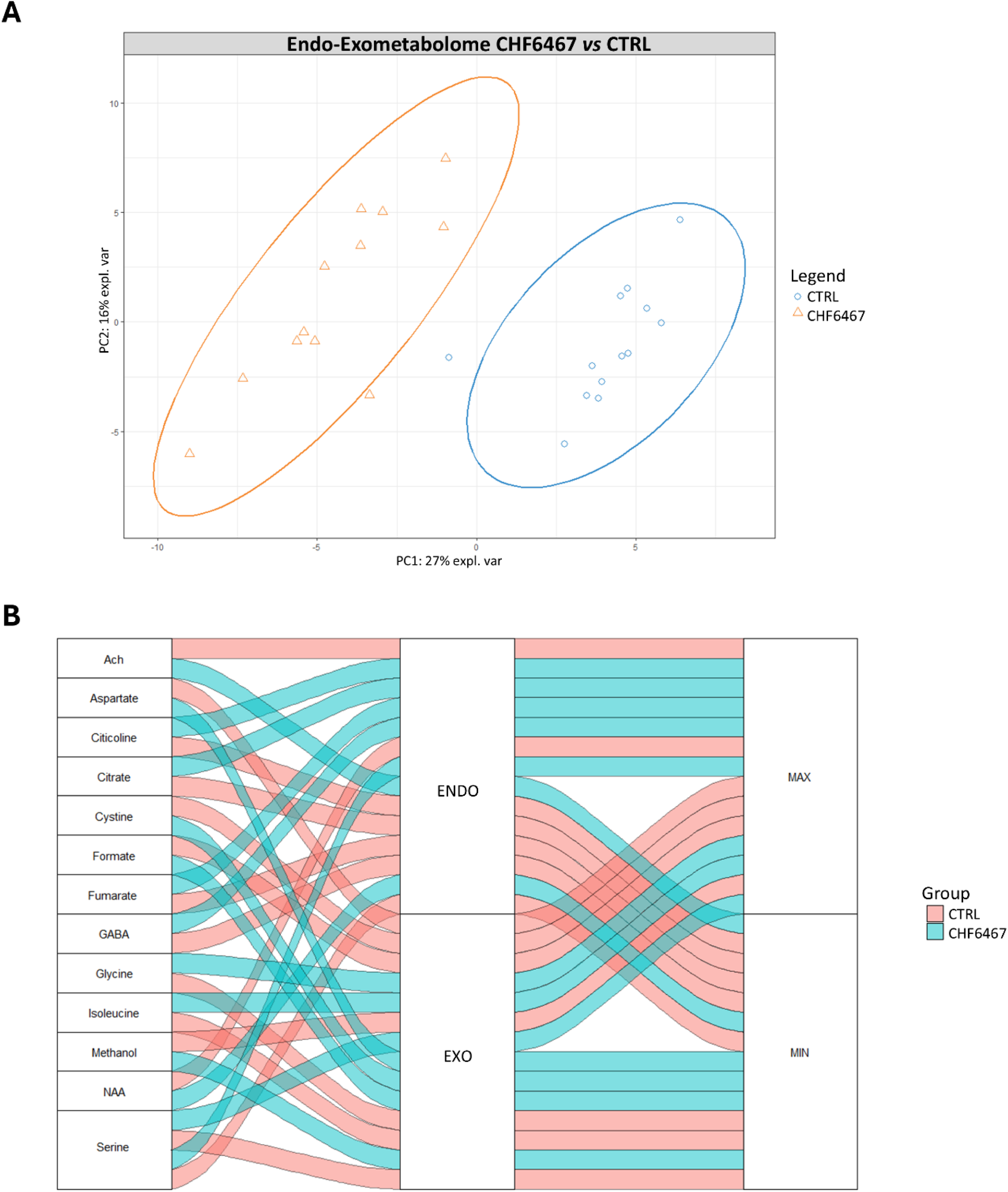
(**A**) sPLS-DA score scatter plots illustrating the metabolomic profiles of combined cellular compartments in SH-SY5Y cells treated with CHF6467 for 24 hours (CHF6467) versus control cells (CTRL). Cluster analyses in the Cartesian space are based on principal components PC1:27% and PC2:18%. sPLS-DA was validated using cross-validation (CV). Results show significant separation of clusters, with accuracy values of 0.96 and 1.0 on PC1 and PC2, respectively, and positive Q2 indices of 0.94 and 0.85, respectively. (**B**) Alluvial plot displaying metabolites that differentiate the clusters identified by sPLS-DA, categorized by VIP value. The first column lists discriminating metabolites, the second indicates the cellular compartment, and the third shows the concentration changes. Max and Min denote the highest and lowest concentrations between the two clusters. Lines connect each metabolite to its respective compartment and the quantitative change, with green lines indicating increased levels in CHF6467-incubated cells, and pink lines indicating typical control cell variations.

The alluvial plot in **Figure 4B** illustrates the metabolites responsible for this separation, highlighting those with VIP scores>1. CHF6467 treatment raised intracellular levels of citicoline, citrate, fumarate, GABA, and serine, while decreasing acetylcholine and N-acetylaspartate (NAA). In the extracellular space, exposure to CHF6467 lowered aspartate, cystine, and formate concentrations, whereas glycine, isoleucine, and serine levels increased.

To explore the connection between intra- and extracellular metabolites, a Cluster Image Map and a Relevance Network Graph were created, showing fumarate as the most connected metabolite (**Figures 5A, B**). Its intracellular levels increased alongside extracellular serine, isoleucine, glycine, and methylamine, while it decreased in relation to proline, cystine, arginine, and formate.

**Figure 5.**
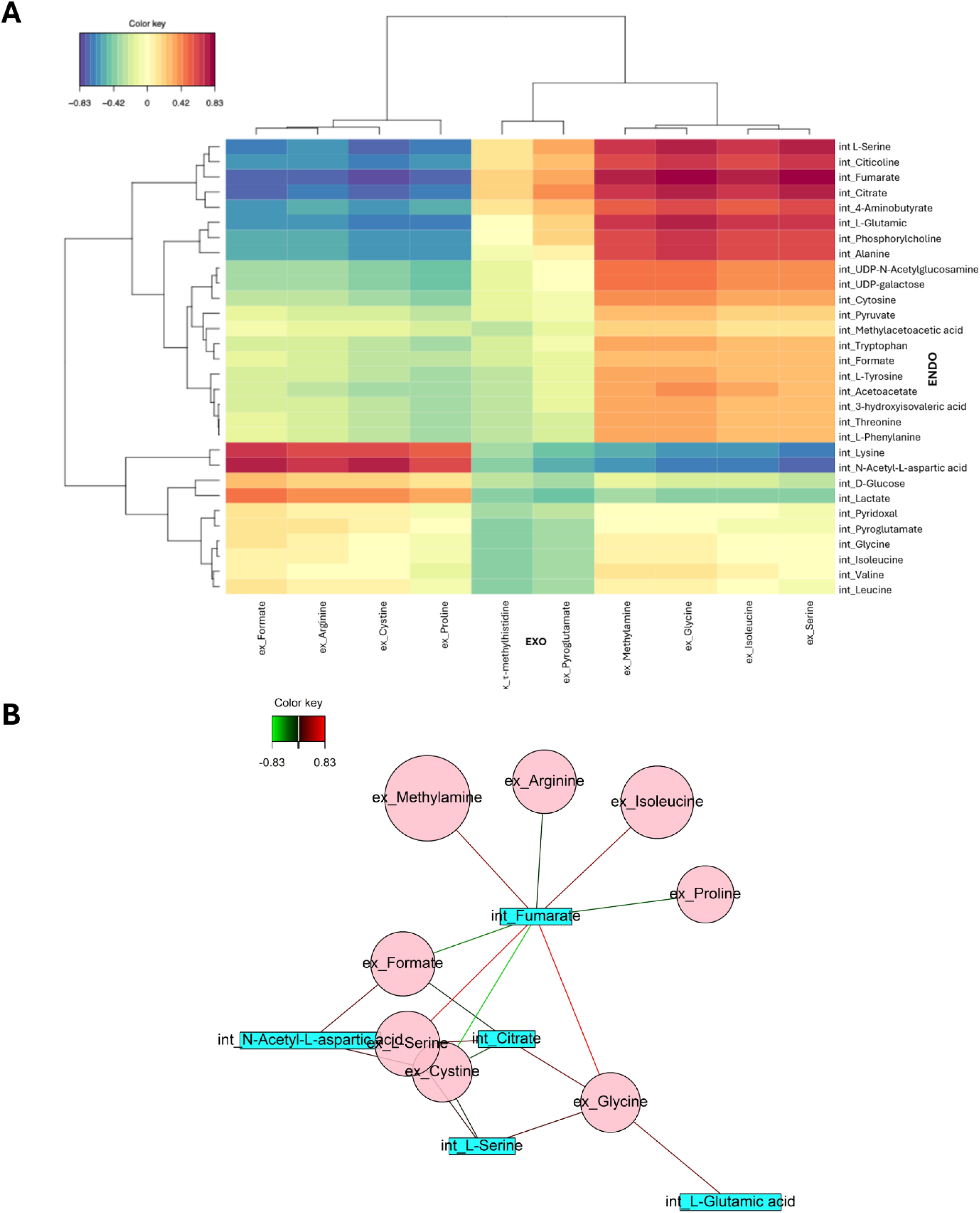
(**A**) Cluster Image Map displaying the relationships between intracellular (columns) and extracellular (rows) metabolites. Pearson’s correlation coefficient was used to analyze linear relationships. Negative correlations are shaded in blue, positive ones in red. (**B**) Relevance Network Graph showing the links between intracellular and extracellular metabolites in SH-SY5Y cells exposed to CHF6467 versus control cells. Extracellular metabolites are depicted as pink circles, intracellular metabolites as blue rectangles. Larger symbols indicate stronger correlations and a stronger influence on cluster separation. Green lines denote indirect correlations, while red lines show direct correlations.

To better understand the biochemical pathways influenced by CHF6467 in relation to the observed metabolite changes, we conducted a pathway enrichment analysis of the intracellular metabolite dataset (**Table 2**).

**Table 2.**
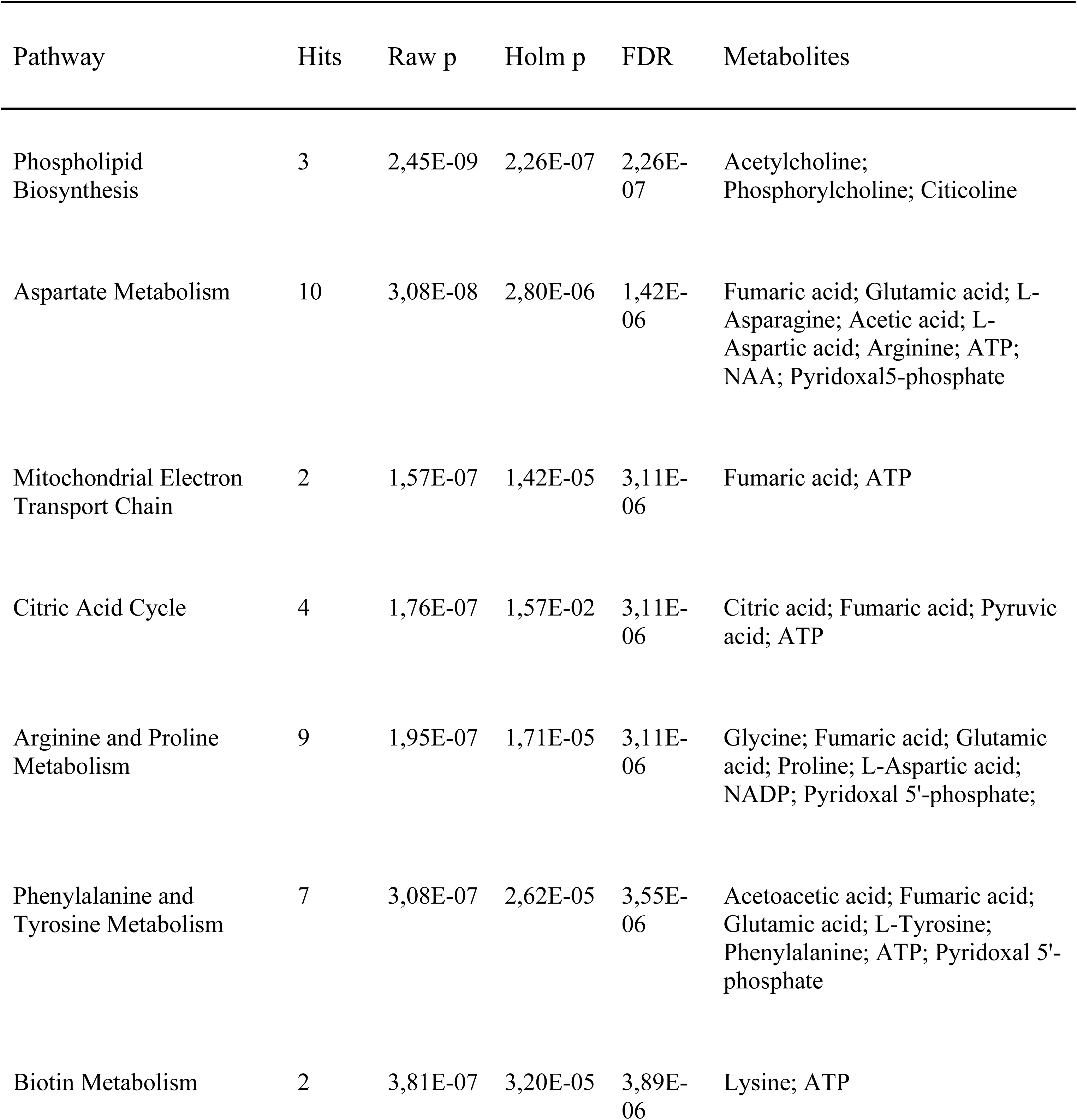
Enrichment pathway analysis: pathway distinguishing CHF6467-treated cells from control cells. For each pathway, the raw p-value, Holm p, FDR value, and involved metabolites are shown. Only pathways with more than one hit and both raw p-values and Holm-adjusted p-values below 0.05 are considered significant.

The most significantly altered pathway was phospholipid biosynthesis, specifically involving phosphatidylethanolamine and phosphatidylcholine. Several amino acid-related pathways were also affected, including arginine and proline metabolism, valine, leucine, and isoleucine degradation, glycine and serine metabolism, selenoamino acid metabolism, and lysine degradation. Changes in neurotransmission were indicated by shifts in the metabolism of aspartate, glutamate, phenylalanine, and tyrosine. Furthermore, pathways related to cellular energy regulation were affected, including the citric acid cycle, electron transport chain, biotin metabolism, propionate metabolism, the urea cycle, and carnitine metabolism.

## Discussion

Alterations in NGF-related cellular processes have been observed in the cholinergic neurons of patients with AD, indicating a role for NGF in disease progression. ^2^ Consistent with these findings, studies in mouse models show that NGF depletion worsens Aβ aggregation and synaptic damage. Conversely, NGF supplementation reduces AD-related neuropathology and improves cognitive function. ^2, 41, 42^ These findings support the idea that NGF may help suppress Aβ-induced neurotoxicity. Additionally, NGF administration has demonstrated therapeutic potential in humans, although it is limited by pharmacokinetic issues and the side effect of pain. ^8, 10–12, 43^ The benefit of the mutant CHF6467 in overcoming side-effect issues motivates further clarification of its mechanism of action.

Pharmacometabolomics has been used as a hypothesis-generating approach in drug discovery to uncover global metabolic perturbations and reveal biological processes warranting further investigation. ^26, 30,44^Consistent with this exploratory perspective, and taking into account the literature data supporting the advantages of the CHF6467 mutant, ^13^ the present study was designed to exploit NMR-metabolomics to initially characterize how insulted neuronal-like cells responds metabolically to the NGF mutant CHF6467, with the identified metabolites and pathways serving as a basis for subsequent functional validation. To this aim, we treated SH-SY5Y neuroblastoma cells with CHF6467 and Aβ(1–42), and profiled intracellular and extracellular metabolites. We preliminarily confirmed CHF6467’s protective effect using a viability assay. Next, we analyzed differences in the metabolomic profile of treated cells, gaining insight into how CHF6467 confers neuroprotection.

Interestingly, our data show that CHF6467 can reverse the metabolic changes caused by Aβ(1–42). This is clearly seen in the sPLS-DA score plot, where the ellipses for control samples and those exposed to both Aβ(1–42) and CHF6467 are close together. Additionally, heatmaps support this, showing that these two groups cluster apart from cells treated only with Aβ(1–42). Pathway enrichment analysis comparing metabolomic profiles of cells treated with Aβ(1–42) alone versus Aβ(1–42) with CHF6467 reveals the biochemical processes influenced by CHF6467 in an *in vitro* system that partially replicates AD changes. Elevated Aβ levels are known to impair neuronal glutamate uptake at synapses, leading to glutamate accumulation outside cells and overstimulation of NMDA receptors. This is also seen in our cell model and aligns with findings in AD models and patient brains. ^45^ Shun et *al.* showed that rat cortical neurons are protected from glutamate toxicity when treated with human or mouse NGF, and observed similar protection against NMDA- and kainate-induced damage. ^38^ In line with these findings, our results indicate that CHF6467 can protect against glutamate-induced neurotoxicity by regulating the glutamate pathway and reducing glutamate levels. Additionally, the decline in glutamate is associated with reduced glycine release, a co-agonist required for full NMDA receptor activation. This combined effect supports the idea that CHF6467 may mitigate excitotoxicity by modulating key components of glutamatergic signaling. Further experiments – such as direct measurements of glutamate levels - are needed to corroborate this hypothesis.

Effects of CHF6467 on the metabolism of arginine and proline are evident, in agreement with their abnormally low concentrationsArginine is involved in several metabolic processes: it serves as a precursor for nitric oxide (NO) synthesis, which can be neurotoxic or neuroprotective in neurons; ^46^ moreover, changes in arginine concentration may affect the metabolism of spermine and spermidine, for which arginine is a precursor. ^47^

CHF6467 appears to influence phospholipids metabolism and the production of their precursors in the presence of amyloid-β peptide. This suggests CHF6467 might help maintain membrane homeostasis, often disrupted in neurodegenerative diseases like Alzheimer’s. Notably, the metabolic shifts caused by CHF6467 are consistent with the maintenance of membrane integrity and suggest a reduction in transmembrane transport impairments associated with amyloid toxicity and neurodegeneration.

As previously demonstrated, exposing SH-SY5Y cells to β-amyloid peptide causes hypoglycemia and elevated 3-hydroxybutyrate levels.^30^ This likely reflects a compensatory response to bioenergetic stress and mitochondrial problems, as well as an attempt to reduce oxidative damage. Notably, 3- hydroxybutyrate is known for its antioxidant properties, scavenging reactive oxygen species (ROS) and influencing redox-sensitive signaling pathways. ^48, 49^ The simultaneous rise in glutathione levels further indicates oxidative stress, probably representing an adaptive increase in the cell’s antioxidant defenses in response to Aβ(1-42). Interestingly, this metabolic response suggests that cells might actively counteract amyloid toxicity by switching to alternative energy sources and boosting redox buffering. When cells are pretreated with CHF6467 before Aβ exposure, there is a notable decrease in ketone bodies and glutathione levels. This suggests a protective role for CHF6467, possibly preventing oxidative stress and reducing the need for compensatory antioxidant responses and alternative energy sources. The rise in pyruvate levels with CHF6467 further supports this idea, potentially reflecting a recovery of glycolytic activity and improved mitochondrial function.

CHF6467 also influenced glucose-related metabolic processes, including glycolysis, gluconeogenesis, the glucose–alanine cycle, and alanine metabolism. This regulation suggests that CHF6467 not only opposes amyloid-induced metabolic reprogramming but also supports a more balanced and efficient energy system. Overall, these results underscore CHF6467’s potential as an early metabolic regulator that helps maintain neuronal homeostasis and reduce the bioenergetic and oxidative issues often linked to neurodegeneration.

To explore how CHF6467 protects against amyloid-β (Aβ) toxicity through metabolic effects, we examined its activity during the pretreatment phase. Specifically, we compared the metabolomic profiles of SH-SY5Y neuroblastoma cells exposed to CHF6467 (100 ng/mL for 24 hours) with those of untreated cells, aiming to identify early metabolic changes that may contribute to CHF6467’s neuroprotective effects. The Relevance Network Graph highlights fumarate as a key component in the CHF6467-induced metabolic landscape. The alluvial plot reveals a notable increase in tricarboxylic acid (TCA) cycle intermediates, such as citrate and fumarate, a finding supported by pathway enrichment analysis. This pattern suggests upregulation of cellular energy metabolism, often associated with conditions that demand higher energy production. Consistent with this, our data indicate modulation of the electron transport chain, where reduced coenzymes from the Krebs cycle support oxidative phosphorylation and ATP synthesis. These results align with previous research in PC12 cells, which showed that NGF enhances glucose metabolism, especially by increasing flux through the TCA cycle. In fact, NGF caused a two-fold rise in pyruvate levels and TCA cycle intermediates in pheochromocytoma cells, indicating a shift toward mitochondrial oxidative metabolism. ^50^

Fatty acid metabolism is also enhanced, as indicated by higher carnitine synthesis. Carnitine is a vital biomolecule needed to transport long-chain fatty acyl-CoAs into mitochondria, where they undergo β-oxidation. Recently, L-carnitine and its acetylated forms have attracted attention as dietary supplements due to their neuroprotective effects. These compounds have shown promise in various conditions involving nerve injury, such as hypoxia-ischemia and AD. ^51^ Therefore, modulating this pathway may indicate an increased energy demand—aligned with previously noted changes in sugar metabolism—and a cellular effort to promote neuroprotection. Overall, these molecular changes suggest a higher cellular energy requirement. Based on these findings, we can infer that CHF6467 provides neuroprotection against Aβ(1-42) toxicity, at least in part by modulating energy-related metabolic pathways during pretreatment. The rise in citicoline, along with lower acetylcholine levels, confirms CHF6467’s ability to influence phospholipid metabolism even before exposure to Aβ(1-42). This shift indicates that choline is being diverted from acetylcholine production toward the synthesis of membrane components, which may improve membrane stability and support structural remodeling.

Consistent with this, pathway analysis indicated that CHF6467 affects the biosynthesis of phosphatidylethanolamine and phosphatidylcholine. These processes are probably important for CHF6467’s involvement in neurite outgrowth, as extending neuronal projections requires significant membrane production. Therefore, CHF6467 might promote increased phospholipid biosynthesis to supply the structural components required for neurite extension. These mechanisms likely help prepare cells to counteract the membrane-disrupting effects of amyloid-β. Additionally, CHF6467 affects molecules involved in neurotransmission, such as glutamate, aspartate, glycine, GABA, and serine—all key mediators of glutamate receptor signaling. After treatment, intracellular GABA levels decrease notably, accompanied by changes in glutamate metabolism. These findings support the idea that CHF6467 indirectly influences the production and/or release of neurotransmitters. Importantly, a close relationship between brain neurotrophic factors and the glutamate–GABA system was proposed as early as 1991. ^52^

Our findings provide novel insights into the mechanisms by which CHF6467 mitigates amyloid-induced metabolic disruption. This is particularly significant given the absence of published metabolomics data regarding NGF or its mutants to date. However, several limitations warrant consideration. First, our metabolomic analysis does not clarify whether the metabolic alterations observed are specifically attributable to CHF6467 or arise from the general activation of the high-affinity TrkA receptor. Second, the use of SH-SY5Y cells—a neuroblastoma-derived cell line— provides an immature neuronal model that may not fully reflect the metabolic nuances of mature, post-mitotic neurons. Furthermore, this monoculture system precludes the investigation of astrocyte-neuron crosstalk and neurovascular interactions, which are integral to in vivo neuronal health’s. ^53^ Despite these constraints, the SH-SY5Y model is recognized as a scientifically rigorous and practical platform for initial mechanistic discovery, circumventing the immediate ethical and technical barriers associated with primary human cultures. ^36^

Overall, **this study** emphasizes the advantage of a pharmacometabolomic approach in exploring its capacity to investigate the molecular mechanism of action of biologically active molecules in an unbiased manner and therefore to uncover unexpected biological outcomes that can further support the drug development process.

## Conclusion

In summary, our study confirms that CHF6467 provides a protective effect against toxicity caused by Aβ(1-42) in neuron-like cells, functioning through the regulation of several vital biochemical pathways:

1. CHF6467 enhances the production of key intermediates necessary for phospholipid synthesis, which are essential for cell membrane structure. This is especially important because membrane integrity is often compromised in neurodegenerative diseases, partly due to amyloid peptide toxicity.
2. CHF6467 manages cellular energy needs and improves the use of bioenergetic substrates, helping to maintain metabolic balance during stress.
3. CHF6467 regulates neurotransmitter levels by influencing glutamate metabolism, which is associated with excitotoxic effects in neurodegenerative diseases.

These results highlight the complex role of CHF6467 in supporting neuronal survival and corroborate its potential as new therapeutic agent for neurodegenerative diseases.

## Material and methods

Dulbecco’s Modified Eagle’s Medium (DMEM), L-glutamine, penicillin and streptomycin, fetal bovine serum (FBS), and CCK-8 were obtained from Sigma-Aldrich (St. Louis, MO, USA).

Aβ(1–42) peptide was expressed in *E. coli* and purified as previously reported.^54^ Following lyophilization, the peptide was subjected to a TFA treatment, a well-established disaggregation protocol that reproducibly generates monomeric Aβ(1–42).^55^ The absence of appreciable β-sheet aggregates in the resulting preparation was verified by ThT fluorescence assay. Aβ(1–42) in its monomeric form was used for all experiments after dilution in DMEM to the desired concentration.

The human neuroblastoma cell line SH-SY5Y was obtained from the American Type Culture Collection (ATCC, Rockville, MD, USA). The cells were cultured in Dulbecco’s Modified Eagle Medium (DMEM, 4500 mg/mL glucose) supplemented with 10% (v/v) fetal bovine serum (FBS), 2 mM L-glutamine, 100 U/mL penicillin, and 0.1 mg/mL streptomycin. The cultures were kept in a humidified incubator at 37 °C with a 5% CO₂ atmosphere and were passaged once they reached confluence.

### Expression and purification of CHF6467

The protocol for expressing and purifying CHF6467 was adapted from the original industrial method provided by Chiesi Farmaceutici. It was scaled down and modified for laboratory use. Specifically, CHF6467 was produced as a proNGF form, ending up in inclusion bodies (IB) within the cytoplasm of E. coli strain BL21 Rosetta^TM^ (DE3). The bacterial strains, transformed with the pET11a- hproNGFP61SR100E plasmid supplied by Chiesi Farmaceutici, were cultivated in LB medium. The medium was also supplemented with ampicillin (100 μg/mL) and chloramphenicol (30 μg/mL). When the cell culture reached an OD600 of 0.6 at 37 °C, gene expression was triggered by adding 1 mM isopropyl-D-thiogalactoside (IPTG). Six hours after induction, cells were harvested and centrifuged to remove the supernatant. The collected cells, containing IBs with the proNGF form of CHF6467, were resuspended in 50 mM Tris and 1 mM EDTA (pH 7.0) and then homogenized. The pellet was weighed and then resuspended in a lysis buffer containing 6 M Guanidine hydrochloride, 2 M urea, 1 mM EDTA, and 50 mM Tris (pH 8.0), using 2.5 mL of buffer per gram of IBs. Dithiothreitol (DTT) was added to the resuspended IB to reach a final concentration of 50 mM, and the mixture was stirred overnight to reduce the disulfide bonds in the proNGF form of CHF6467. After reduction, the pH was lowered to 3.5 with HCl, and the solution was filtered through a 0.65 µm positively charged glass fiber filter (Sartopure® GF Plus Sartoscale 25, Sartorius, Germany). DTT was then removed by constant volume diafiltration with a solution containing 50 mM Tris, 2 M urea, 6 M Guanidine hydrochloride, and 1 mM EDTA at pH 3.5, followed by concentrating the solution. The buffer exchange used seven times the concentrate volume, and all diafiltration and concentration steps employed an Amicon® Stirred Cell (Merck-Millipore, Germany) with 10 kDa ultrafiltration discs made of polyethersulfone (Biomax, Italy).

To refold the proNGF form of CHF6467, the solution was added to the refolding buffer (50 mM Tris, 1 M L-arginine HCl, 0.1 M NaCl, 0.35 M NaOH, 5 mM EDTA, 5 mM GSH, 1 mM GSSG, pH 9.5) at a 1:50 ratio of protein solution to buffer. It was dispensed using a syringe pump at a steady flow rate of 2 μL/min at 4°C. During addition, the buffer was continuously stirred. After complete addition, stirring continued for 8 hours. Then, the pH was adjusted to 7.5 with HCl, and the solution was concentrated. Buffer exchange was performed using 1 M arginine HCl and 20 mM sodium phosphate (pH 7.4) via constant volume diafiltration against concentrated refolding buffer. The retentate was diluted with 10 mM sodium phosphate (pH 7.3) until conductivity was below 25 mS/cm. Finally, to eliminate any precipitate, the solution was filtered through a 0.65 µm positively charged glass fiber filter.

Before trypsin cleavage, CaCl_2_ was added to a final concentration of 20 mM, and the pH was adjusted to 7.0 using NaOH. Trypsin was then added to the proNGF form of CHF6467 at a final concentration of 1% (w/w), and the reaction was stirred continuously for 4 hours at 4°C. Analyzing the trypsin time course with SDS-PAGE, the 4-hour duration was identified as most effective for cleavage, since shorter times left higher residual proNGF levels, lowering the yield (data not shown). Trypsin activity was halted by adding PMSF to 1 mM and aprotinin (BPTI) to 2 µg/mL.

After trypsin cleavage, the solution’s pH was adjusted to 5.5–6.0 with HCl, then filtered through a 0.65 µm positively charged glass fiber filter. The purification was performed via ion-exchange chromatography: the CHF6467 solution was loaded onto a CaptoMMC column (Sigma-Aldrich Merck, Germany) at 1 mL/min using an AKTA purifier system. The column was equilibrated with 350 mM Arg-HCl and 20 mM phosphate buffer, pH 6.0. After loading, it was washed with the same buffer. The protein was eluted with 350 mM Arg-HCl, 50 mM phosphate, pH 9.5, resulting in a single peak. The eluate was then dialyzed against phosphate buffer pH 6.0 to reduce arginine levels. The identity of CHF6467 was confirmed via Electro-Spray Mass Spectrometry using a Vanquish Flex UHPLC system coupled with an Orbitrap Exploris 120 and HESI-II source (Thermo Fisher Scientific, Milan, Italy).

### Cell viability test

Cell viability was evaluated by measuring mitochondrial metabolic activity with the Cell Counting Kit-8 (CCK-8, Cat. CK04, Dojindo Laboratories, Rockville, MD, USA), following the manufacturer’s instructions. To test CHF6467’s potential to protect SH-SY5Y cells from Aβ (1-42) toxicity, 8 × 10³ cells per well were plated in 96-well plates for 24 hours, then treated with 100 ng/mL or 200 ng/mL CHF6467 for another 24 hours. Subsequently, 50 μM Aβ (1-42) peptide was added for 48 hours. The CCK-8 reagent was diluted in culture medium (10%) and incubated for 1 hour. Absorbance at 450 nm was measured using a microplate reader (Multiskan Go, Thermo Scientific, Waltham, MA, USA). Cell viability was expressed as a percentage of viable cells relative to the untreated control. Data are shown as mean ± SD from three independent experiments. Statistical analysis involved one-way ANOVA followed by Dunnett’s test for multiple comparisons, conducted with GraphPad Prism 8.0 (GraphPad Software, San Diego, CA, USA). Significance was set at p < 0.05. Symbols indicate significance levels: #### for p < 0.0001 vs. CTRL; *, ** for p < 0.05 and p < 0.01 versus Aβ (1-42).

## 1H NMR Metabolomics

### Exposure of SH-SY5Y Cells to CHF6467 and Aβ(1–42)

To prepare metabolomic samples, cells were cultured in 60 mm dishes and allowed to adhere overnight. For co-treatment, cells were pretreated with CHF6467 (100 ng/mL), followed by the addition of Aβ (1-42) after 24 hours, and then incubated for an additional 48 hours at a sub-toxic concentration (5 μM). Control samples included cells exposed only to Aβ (1-42) at the same concentration and duration, while the control group was treated solely with the vehicle for the same period.

### Sample Collection and Intracellular Metabolites Extraction

Following treatments, both the culture media and cell pellet were collected for metabolomics analysis, focusing on the exometabolome and endometabolome, respectively. The medium was transferred to microcentrifuge tubes and centrifuged at 1000× g for 10 minutes. The same procedure was performed on cell-free medium incubated under identical conditions. Supernatants were then transferred to new microcentrifuge tubes and stored at −80 °C until NMR analysis. After removing the media, cell dishes were rinsed with cold PBS (pH 7.4) to eliminate media residues, and the cells were scraped in methanol. To extract intracellular metabolites from the cell pellet, homogenization and biphasic extraction were performed using equal volumes of methanol, chloroform, and water. ^56^ Samples were centrifuged at 6000 rpm for 10 minutes at 4 °C, separating polar and non-polar phases. Polar extracts from the cell pellet were dried under vacuum with a SP-Genevac EZ-2 4.0 concentrator, while lipophilic extracts were dried with nitrogen flow for subsequent analysis. All extracts were stored at −80 °C before NMR testing.

### NMR Sample Preparation

Lyophilized cell extracts were reconstituted with 200 μL of buffer (50 mM Na_2_HPO_4_, 1 mM trimethylsilyl propionic-2,2,3,3-d_4_ acid sodium salt (TSP-d_4_), 10% D_2_O). TSP-d_4_ served as an internal standard for aligning and quantifying NMR signals. For analyzing growth media, 100 μL of cell medium was diluited with 100 μL of the same phosphate buffer. These samples were subsequently transferred to 3 mm NMR tubes for ^1^H NMR analysis.

### NMR Data Acquisition and Processing

1D ^1^H NMR spectra were acquired using a Bruker AscendTM 600 MHz spectrometer equipped with a 5 mm triple resonance Z gradient TXI probe (Bruker Co, Rheinstetten, Germany) at 298 K. One-dimensional NOESY NMR spectra were recorded with 20,000 points, a spectral width of 12 ppm, an acquisition time of 1.36 seconds, a relaxation delay of 5 seconds, a mixing time of 10 ms, and 128 scans. ^57^ Spectrometer control and data analysis were performed using Topspin v3.0 (Bruker Biospin). Spectral analysis adopted an untargeted metabolomics approach, identifying each metabolite before statistical testing with Chenomx NMR-Suite v10.1 (Chenomx NMR suite, v10.1, Edmonton, AB, Canada). Quantification of the 1D NMR data was conducted using NMRProcFlow v1.4.10, and the resulting data matrix was subsequently subjected to statistical analyses.

## Statistical analysis

The sample data were normalized utilizing the sum, subsequently subjected to log transformation and Pareto scaling prior to analysis utilizing the MixOmics R package version 6.28.0 (mixOmics) and MetaboAnalyst 6.0, as cited in reference. ^58, 59^ Univariate analysis was independently performed on the exometabolome and endometabolome segments of the groups, employing T-Test and Fold Change metrics, with the outcomes illustrated in a Volcano plot. ^60^ To enhance data accuracy and facilitate the extraction of biological insights, multivariate statistical analysis (MVA) was initially applied to the concentration matrices of both the exometabolome and endometabolome, followed by an analysis of the combined dataset.

To analyze both the endo- and exometabolome, we used the mixOmics R package with a P-integration approach. To improve accuracy and reduce the impact of inherent variability between matrices, we further removed batch effects using the R limma package. ^61^ Multivariate analysis (MVA) was performed on the combined matrices of endo- and exo-metabolites using supervised sparse Partial Least Squares (sPLS), also called the projection to latent space method. ^62^

The sPLS analysis employed an LASSO penalty on the loading vectors to minimize the number of original variables used in creating the latent variables. ^63^ A sample plot demonstrates how samples’ metabolomic profiles cluster, with each point representing a sample based on its projection onto the selected latent components. Leave- one- out cross- validation validated the model using R 2, Q 2, and accuracy metrics.^ 64^ Additionally, sPLS models were validated through ROC curve analysis. ^65^

Variable correlations are visualized via a circular correlation plot, where all vectors are placed within a unit circle (radius 1). The position of each vector indicates its correlation with the components- longer vectors signify stronger associations. Vectors close to each other indicate high correlation. ^58^ The contribution of each variable is displayed in a bar graph, with color- coding to highlight the maximum value between two clusters, showing where the metabolite’s concentration is highest. To depict metabolite changes quantitatively, heatmaps were created using normalized data, average group concentrations, and Euclidean distance. ^66^ Intra- and extracellular metabolite correlations were analyzed using a Cluster Image Map built on the Pearson correlation coefficient. This coefficient measures the strength and direction of relationships between two variables, ranging from –1 to + 1. Values near- 1 indicate inverse correlation, while those near + 1 indicate direct correlation, with significance increasing towards the extremes. ^67^

The Relevance Network Graph includes metabolite centrality, indicating how much individual metabolites contribute to distinguishing metabolic profiles. Only metabolites with a Pearson correlation coefficient above 0. 70 and a high degree- based on the number of variables in the joint matrix of exometabolome and endometabolome- were visualized. The degree reflects the number of connections a node has, serving as a measure of centrality. ^68^ Pathway analysis was performed using The enrichment pathway tool in Metaboanalyst 6. 6.0. KEGG pathways were selected based on low false discovery rates (FDR), p- values less than 0. 05, and at least two hits (number of metabolites involved). ^69^

## Supporting Information

**Figure S1**. Performance evaluations of the sPLS-DA model based on ^1^H-NMR data involved ROC curve analysis of the combined exo and endometabolome profiles from CHF6467_Abeta and Abeta compared to CTRL. The ROC values for the first and second components, along with the corresponding *p-*values, are listed in the table.

**Figure S2.** Performance evaluations of the sPLS-DA model based on ^1^H-NMR data involved ROC curve analysis of the combined exo and endometabolome profiles from CHF6467 versus CTRL. The ROC values for the first and second components, along with the corresponding *p-*values, are listed in the table.

---

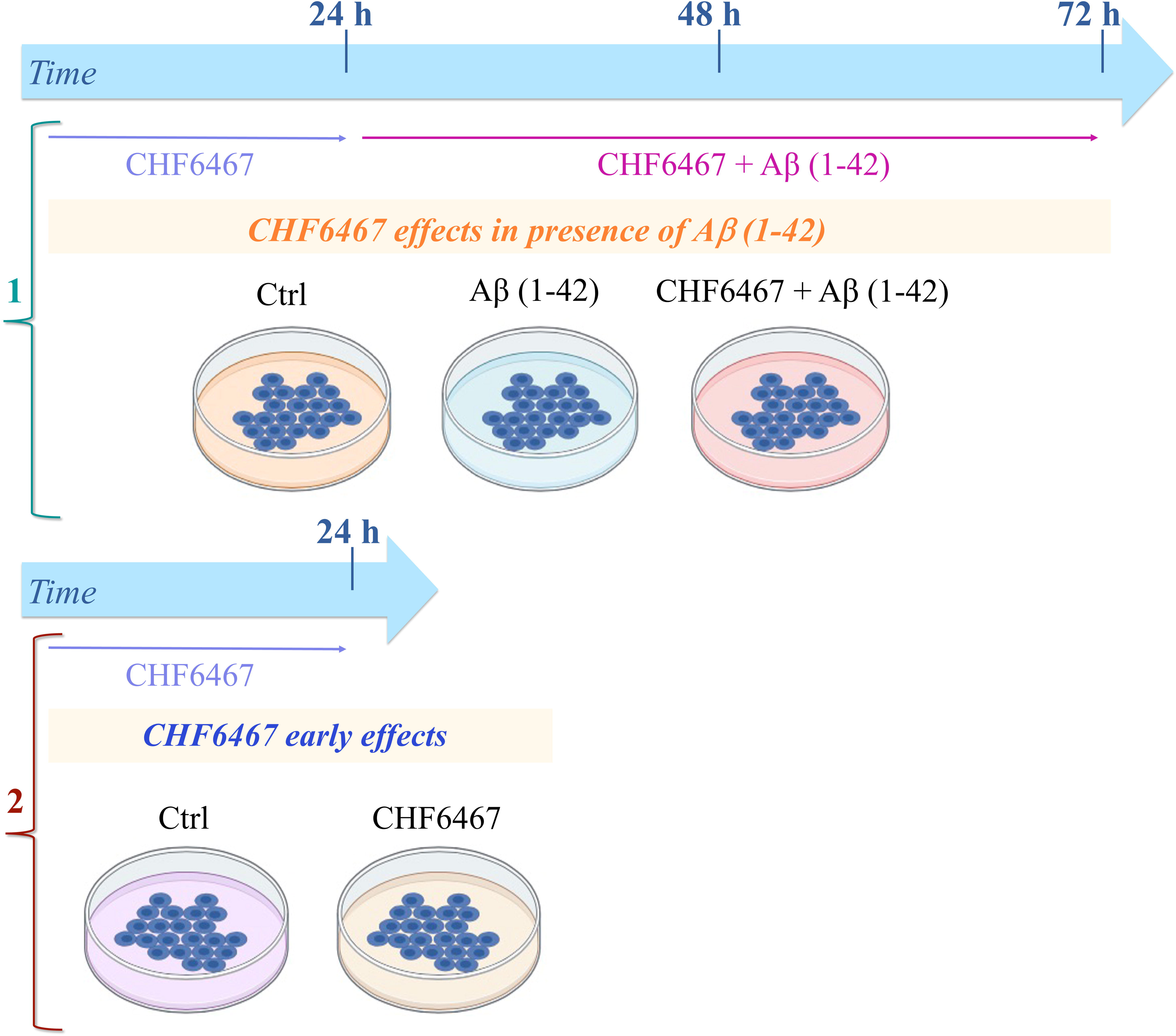

