## Supporting Information for "The protective effect of NGF mutant CHF6467 in SH SY5Y cells exposed to Aβ(1–42): a pharmacometabolomic approach"

**Figure S1.** Performance evaluations of the sPLS-DA model based on $^1$H-NMR data involved ROC curve analysis of the combined exo and endometabolome profiles from CHF6467_Abeta and Abeta compared to CTRL. The ROC values for the first and second components, along with the corresponding *p*-values, are listed in the table.

**Figure S2**. Performance evaluations of the sPLS-DA model based on $^1$H-NMR data involved ROC curve analysis of the combined exo and endometabolome profiles from CHF6467 versus CTRL. The ROC values for the first and second components, along with the corresponding *p*-values, are listed in the table.

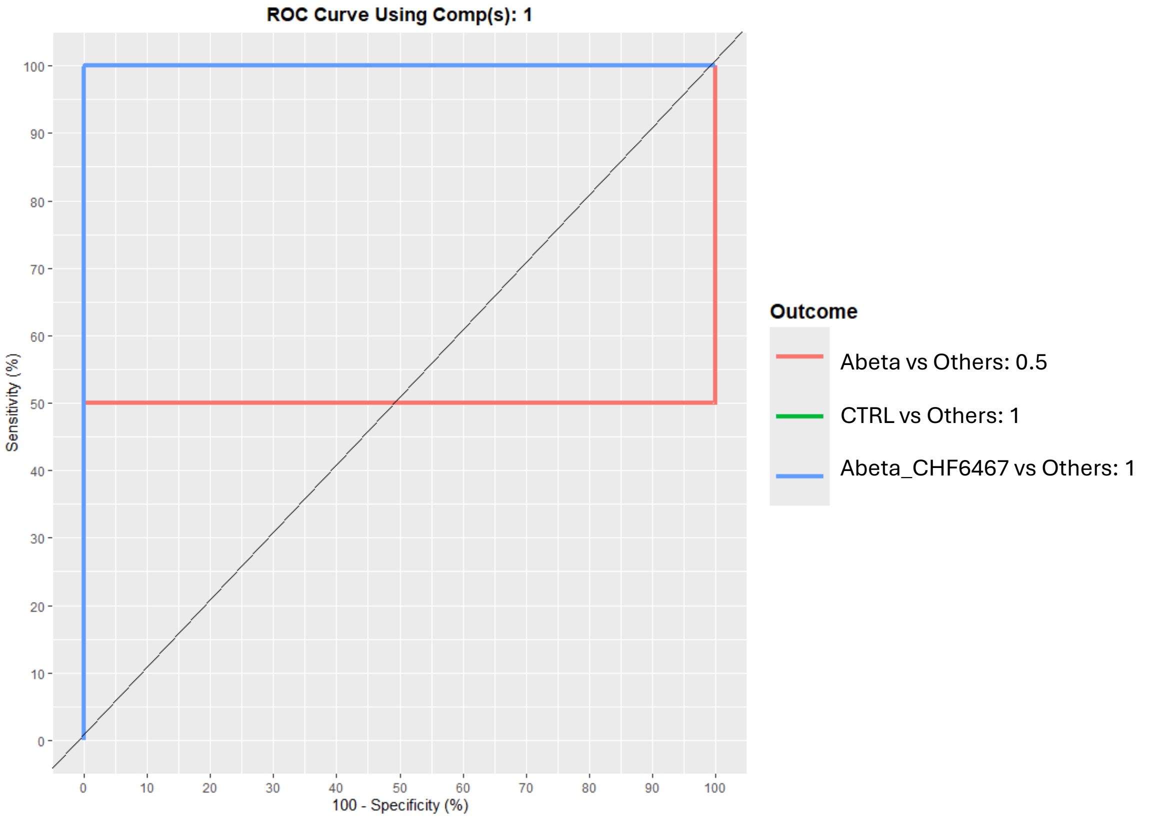


| Outcome | AUC Component 1 | *p-value* |
|---|---|---|
| **Abeta vs Others** | 0.50 | 0.003 |
| **CTRL vs Others** | 1 | 1.25E-11 |
| **Abeta_CHF6467 vs Others** | 1 | 1.48E-09 |

**Figure S1.** Performance evaluations of the sPLS-DA model based on $^{1}$H-NMR data involved ROC curve analysis of the combined exo and endometabolome profiles from CHF6467_Abeta and Abeta compared to CTRL. The ROC values for the first and second components, along with the corresponding *p*-values, are listed in the table.

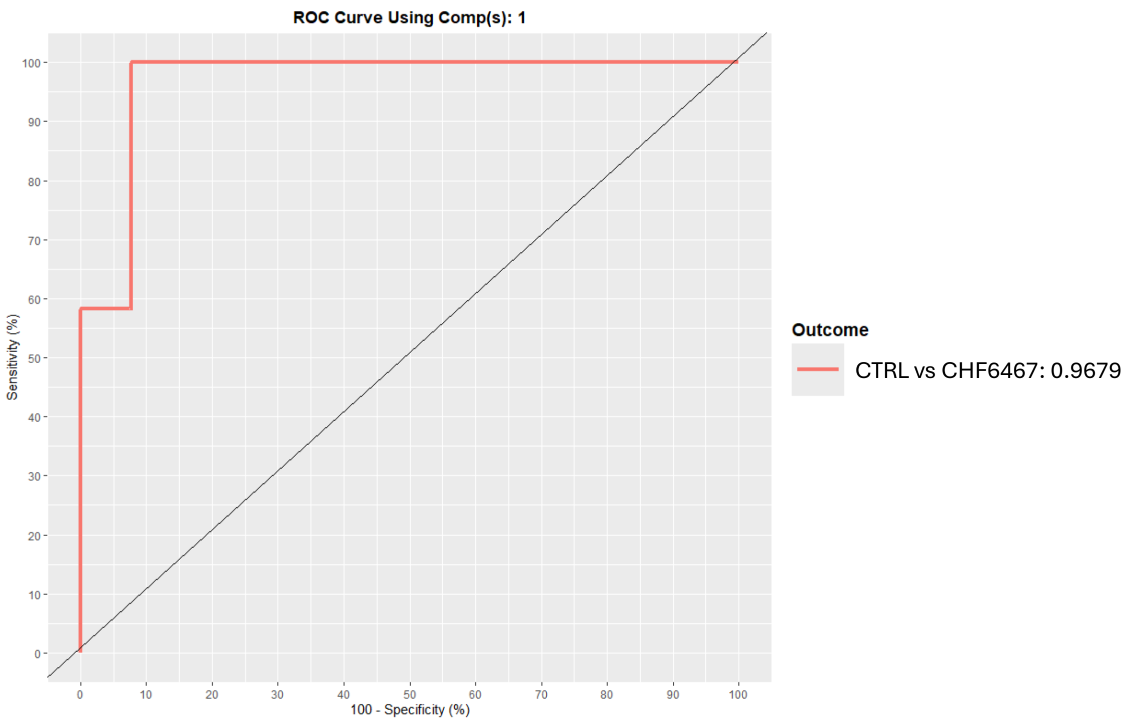


| **AUC Component 1** | ***p-value*** |
|---|---|
| 0.97 | 1.25E-10 |
| **AUC Component 2** | ***p-value*** |
| 0.96 | 1.15E-09 |

**Figure S2.** Performance evaluations of the sPLS-DA model based on $^{1}$H-NMR data involved ROC curve analysis of the combined exo and endometabolome profiles from CHF6467 versus CTRL. The ROC values for the first and second components, along with the corresponding *p*-values, are listed in the table.
